# Transcriptome Dynamics in Mouse Amygdala under Acute and Chronic Stress Revealed by Thiol-labeled RNA Sequencing

**DOI:** 10.1101/2024.01.12.573386

**Authors:** Dan Zhao, Lu Zhang, Yang Yang

**Author notes:** These authors contributed equally to this work (Dan Zhao, Lu Zhang).

## Abstract

Both acute and chronic stress have significant impact on brain functions. The amygdala is essential in mediating stress responses, but how its transcriptomic dynamics change under stress remains elusive. To overcome the difficulties in detecting subtle stress-induced changes by evaluating total RNA using classic RNA sequencing, we conducted thiol-labeled RNA sequencing (SLAM-seq). We injected 4-thiouridine (4sU) into mouse amygdala followed by SLAM-seq to detect nascent mRNA induced by acute and chronic restraint stress, and found that SLAM-seq could label actively transcribed genes in the major neuronal and glial subtypes. We also found that acute stress induced active transcription of 6 gamma-aminobutyric acid (GABA) receptors and only 1 glutamate receptor, indicating an imminent increase of inhibitory control in the stressed amygdala. Conversely, chronic stress led to active transcription of 3 glutamate receptors and 4 GABA receptors, suggesting a release of inhibitory control and hyperactivity of the amygdala. SLAM-seq also identified genes associated with myelination under chronic stress, and this finding is confirmed by immunostaining showing increased myelination in chronically stressed amygdala. Additionally, genes detected by SLAM-seq and RNA-seq only partially overlapped, with SLAM-seq particularly sensitive to transcriptional changes in genes with high basal transcription levels. Thus, by applying SLAM-seq *in vivo*, we obtained a rich dataset of nascently transcribed genes in the amygdala under stress, and revealed distinct transcriptional dynamics associated with acute and chronic stress.

## Introduction

The impact of stress on the brain is profound and has been linked to an increased risk of mental illnesses, including depression and anxiety disorders [1, 2]. Both acute and chronic stress can induce adaptive changes in various brain regions [3–9], including the amygdala, which plays a key role in processing physiological and behavioral responses to stress [10, 11]. In its resting state, the amygdala is tightly regulated through inhibitory mechanisms mediated by gamma-aminobutyric acid (GABA) [12]. Previous research showed that stress leads to amygdala disinhibition, resulting in its heightened activity [13, 14]. While many studies have demonstrated that stress affects amygdala function, the underlying molecular mechanisms responsible for these changes remain elusive. To understand the stress-induced effects in the amygdala at the molecular level, it is essential to study how gene expression in amygdala is influenced by acute and chronic stress.

It has long been established that neuronal activities lead to fast expression of immediate early genes, which further regulates transcription of thousands of downstream genes over an extended period [15–17]. Such activity-dependent transcription dynamics are crucial for modulating neuronal functions and long-term connectivity changes [18, 19]. Studies have shown that glial cells in the brain also exhibit significant functional changes induced by neuronal activities. For example, oligodendrocytes undergo activity-dependent myelination [20, 21]. However, the abundance of gene transcripts for maintaining cellular functions, and the complex morphology of neurons and glia, made the classic RNA sequencing (RNA-seq) method unsuitable for studying the dynamic transcriptomic changes of these cells [22].

The thiol-labeled RNA sequencing (SLAM-seq) method specifically label newly transcribed mRNAs [23], and has been widely applied in cultured cells [24, 25]. Recently, SLAM-seq has also been extended to *in vivo* studies to explore genomic changes during zebrafish embryo development [26], and to identify cell type-specific transcriptome in mouse tissues in a Cre-dependent manner (SLAM-ITseq, [27]). Although SLAM-ITseq provides a solution to metabolically label RNA with 4-thiouracil (4-TU) in a specific cell type *in vivo*, this method requires usage of double-transgenic mouse lines to achieve cell type-specific expression of uracil phosphoribosyltransferase (UPRT), to transform 4-TU to 4-thiouridine (4sU) in Cre-expressing cells, limiting the applicability.

By conducting SLAM-seq with direct stereotactic injection of 4sU, we detected newly transcribed genes in the amygdala under acute and chronic stress. Bioinformatic analyses showed that the injected 4sU was integrated into the transcription of genes expressed in major brain cell types. In addition, acute stress and chronic stress induced differential transcription of GABA and glutamate receptors, indicating diverse inhibitory control in the amygdala under acute and chronic stress. Furthermore, compared to RNA-seq, SLAM-seq is more sensitive to transcriptional changes in genes expressed in high basal levels.

## Results

### Adapt SLAM-seq for the brain through stereotactic injection of 4sU

Neuronal activity is known to quickly trigger the transcription of many genes [15–17]. Seeking to detect gene transcription in response to diverse conditions of neuronal activity in different brain regions, we adapted the SLAM-seq protocol *in vivo* by injecting 4sU directly into the brain region of interest. The stereotactic injection method was used to ensure the target delivery of 4sU, a nucleotide analog that can be imported into animal cells by equilibrate nucleoside transporters [28], in selected brain regions (See Methods for details). To estimate the 4sU incorporation rate in brain cells, we first injected 4sU (100mM, dissolved in 10% DMSO, 1uL) into the mouse auditory cortex (**Fig. 1a**), and performed biotin dot blot essay with mRNAs extracted from the cortex at 2 h, 8 h, and 24 h after 4sU injection [29]. The labeling strength increased with labeling time (**Fig. 1b**), and at 24 h post injection, the labeling is highly efficient. Thus, we determined that 24 h is a sufficient duration to allow 4sU diffusion into brain cells to support SLAM-seq.

**Fig. 1:**
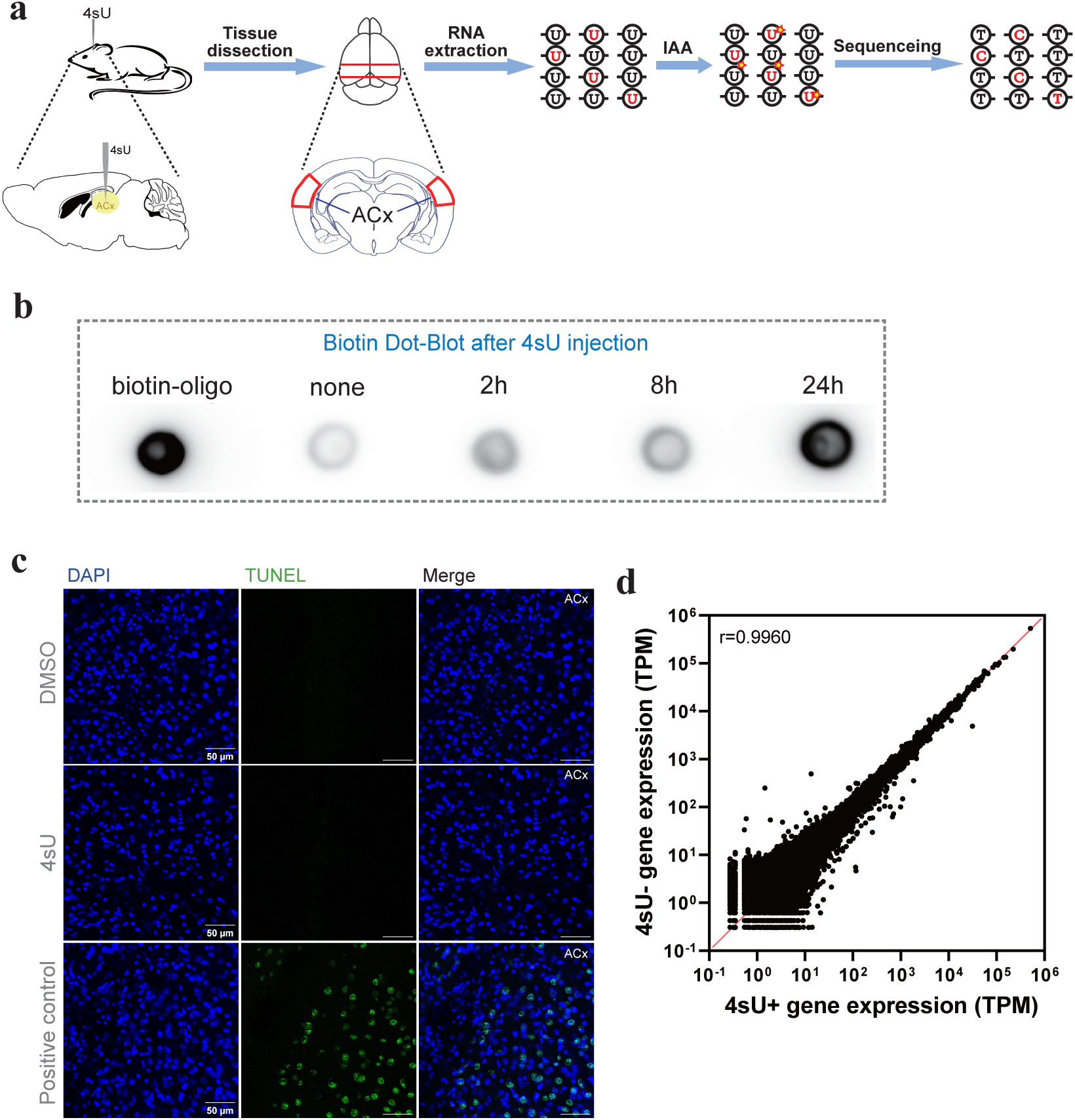
SLAM-seq for mouse brain cells by stereotactic injection of 4sU into the brain. **(a)** Schematic showing SLAM-seq procedure, including stereotactic 4sU injection, RNA extraction, iodoacetamide (IAA) treatment, RNA sequencing and T-C conversion detection. **(b)** Dot blot essay of RNA extracted from mouse brain labeled with 4sU for different durations. Left to right: biotin-oligo (positive control); RNA samples extracted from mouse auditory cortex with no 4sU injection (negative control); 2 h, 8 h, and 24 h after 4sU injection. **(c)** TUNEL staining of brain slices for positive control (top panels), and brain slices with DMSO (middle) and 4sU (bottom) injections in auditory cortex. Green signals denoted dead cells. Scale bar, 50 μm. **(d)** Correlation of RNA levels extracted from DMSO (denoted as 4sU-) and 4sU (denoted as 4sU+) injected auditory cortex samples. ρ, spearman correlation coefficient.

To evaluate the extent of tissue damage potentially caused by the injection of 4sU, we used TUNEL (Terminal deoxynucleotidyl transferase dUTP Nick End Labeling) staining to assess cell death in the injected brain region [30]. We injected 4sU (100mM, dissolved in 10% DMSO, 1uL) and DMSO (10%, 1uL) into the mouse auditory cortex and amygdala, and 24 h later, extracted the mouse brains for slicing and TUNEL staining (Methods). We did not see dead cells in the brain slices near the injection sites, for both 4sU and DMSO injections, suggesting that neither 4sU nor DMSO induced detectable cell death or tissue damage in the injected brain regions (**Fig. 1c, Supp.** Fig. 1).

We also assessed whether the presence of 4sU had any impact on the transcriptome by performing 150bp paired-end sequencing of total mRNA extracted from the isolated mouse auditory cortex and amygdala at 24 h after stereotactic 4sU or DMSO injections (**Supp.** Fig. 2a**, 2b**). By comparing the read counts for all genes [23], we did not observe any significant difference in gene expression levels between 4sU (denoted as 4sU+) and DMSO (denoted as 4sU-) groups (**Fig.1d, Supp.** Fig. 2c). These findings support that local injection of 4sU into specific brain regions enables efficient incorporation into transcription in brain cells, without obviously affecting the gene expression.

### Identify 4sU-incorporated mRNA with SLAM-seq under basal conditions

Recently, in addition to cultured cells [24, 25], SLAM-seq has also been applied *in vivo* by injecting 4sU into zebrafish embryo [26], or injecting 4-thiouracil (4-TU) intraperitoneally into transgenic mice expressing UPRT in selected cell types [27]. Given that local injection of 4sU into the mouse brain has never been reported, we needed to empirically determine whether the delivered 4sU can be incorporated into the nascently transcribed RNA molecules. We injected 4sU (100mM, 1uL, dissolved in 10% DMSO) and DMSO (10%, 1uL) into the mouse amygdala 24 h prior to tissue dissection and RNA extraction (**Fig. 2a**), to establish a baseline (BL) for nascent RNA transcription in normal behaving mice. We then subjected the aforementioned total RNA extracts to iodoacetamide (IAA) treatment (Methods, **Supp.** Fig. 3a), to covalently attach a carboxyamidomethyl group to 4sU by nucleophilic substitution [23], and then performed 150bp paired-end RNA-seq (**Supp.** Fig. 3b). During the reverse transcription step of RNA-seq library preparation, a guanine (G), instead of an adenine (A), is base paired to an alkylated 4sU, leading to the thymine (T) to cytosine (C) base conversion (T>C conversion) at the corresponding T position in mRNA [23]. To control for the background T>C conversion rate without 4sU, RNA obtained from DMSO-injected amygdala was subjected to the same procedures before RNA-seq.

**Fig. 2:**
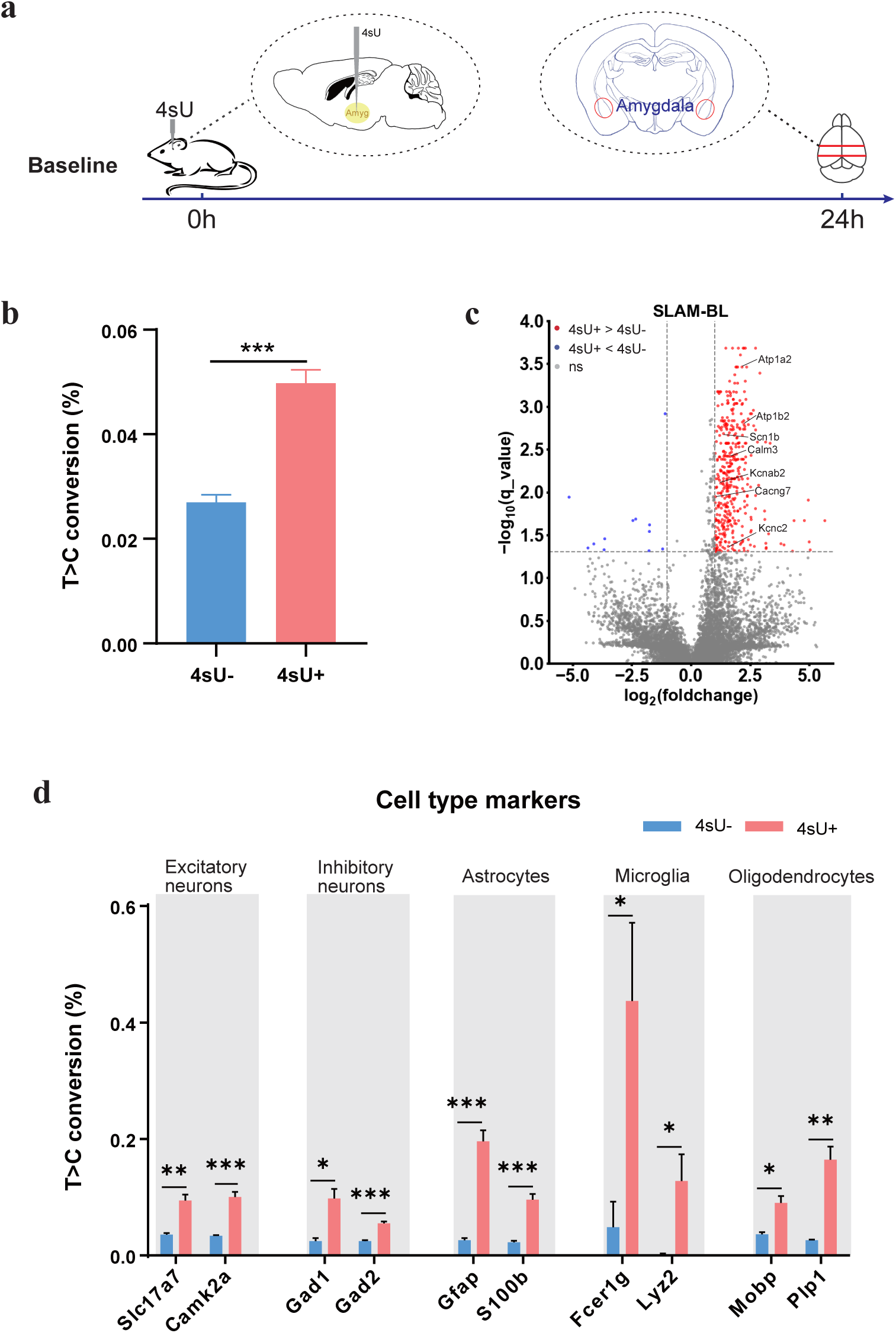
Identifying 4sU-labeled genes with SLAM-seq and match to different cell types. (a) Time line showing stereotactic 4sU injection and brain extraction under baseline (BL) condition. Shading demonstrates nascent mRNA that can be detected by SLAM-seq. (b) T>C Conversion rates of total mRNA extracted from DMSO (n=5) and 4sU (n=4) injected amygdala under baseline condition. (c) Volcano plots showing T>C conversion rates of transcripts under baseline, with or without 4sU injection. Red dots represent transcripts with significant higher T>C conversion rates in the 4sU+ group (FDR < 0.05 and log_2_FC > 1); blue dots represent transcripts with significant lower T>C conversion rates in the 4sU+ group (FDR < 0.05 and log_2_FC < -1). (d) T>C Conversion rates of marker genes for multiple neuron and glia cell types.

We calculated the T>C conversion rates for each gene in both the 4sU-injected and DMSO-injected samples using SLAM-DUNK software [31]. At the whole transcriptome level, the average T>C conversion rate of the 4sU-injected group was significantly higher than that of DMSO-injected group (**Fig. 2b**), indicating that the injected 4sU was incorporated into transcription. We identified 414 genes with significantly higher T>C conversion rates in the 4sU group than in the DMSO group (**Fig. 2c**), which can be plausibly interpreted as genes undergoing active transcription. Note that these mice were not subjected to any behavioral or physiological perturbations, therefore, the 414 differentially 4sU-labeled genes were actively transcribed under normal baseline conditions. Indeed, these genes included sodium channels (*Scn1b*), potassium channels (*Kcnab2, Kcnc2*), cation transport ATPases (*Atp1a2, Atp1b2)*, calcium channels (*Cacng7*) and Calmodulin (*Calm3*), all of which are essential for maintaining basic neuronal functions [32].

### Determine the 4sU labeling efficiency in distinct cell types

Identification of genes encoding neuron-specific ion channels indicated that 4sU was successfully incorporated into transcription of genes in the amygdala neurons. To additionally evaluate the efficiency of 4sU incorporation into the transcription of other cell types in amygdala, we cross-referenced the 414 differentially 4sU-labeled genes with known marker genes of major cell types as reported in single-cell RNA sequencing studies [33, 34]. We found that the 414 differentially labeled genes under baseline conditions covered marker genes of excitatory (*Slc17a7, Camk2a*) and inhibitory neurons (*Gad1, Gad2*), astrocytes (*Gfap, S100b*), microglia (*Fcer1g, Lyz2*) and oligodendrocytes (*Mobp, Plp1*). The T>C conversion rates of these genes were significantly higher in 4sU-injected than DMSO-injected amygdala (**Fig. 2d**). These results indicate that 4sU has been imported into major neuronal and glial cell types of the amygdala, and could be effectively incorporated into the transcription of these cells to signal nascent transcription.

### Transcriptome dynamics of the amygdala under acute stress

The amygdala has been extensively implicated in stress responses [10, 11]. To identify genes that were actively transcribed upon acute stress (AS), we subjected 4sU- and DMSO-injected mice to 1.5 h of restraint stress [35], and immediately after the stress treatment, we isolated the amygdala and extracted total RNAs (Methods, **Fig. 3a**). We treated the RNA samples with IAA for SLAM-seq, and identified the differentially 4sU-labeled genes using SLAM-DUNK by comparing T>C conversion rates between 4sU- and DMSO-injected amygdala samples (**Supp.** Fig. 4a). Meanwhile, we conducted a separate bulk RNA-seq analysis, using DESeq2 software package [36] to identify differentially expressed genes (DEGs) in acute stress condition, compared to baseline.

**Fig. 3:**
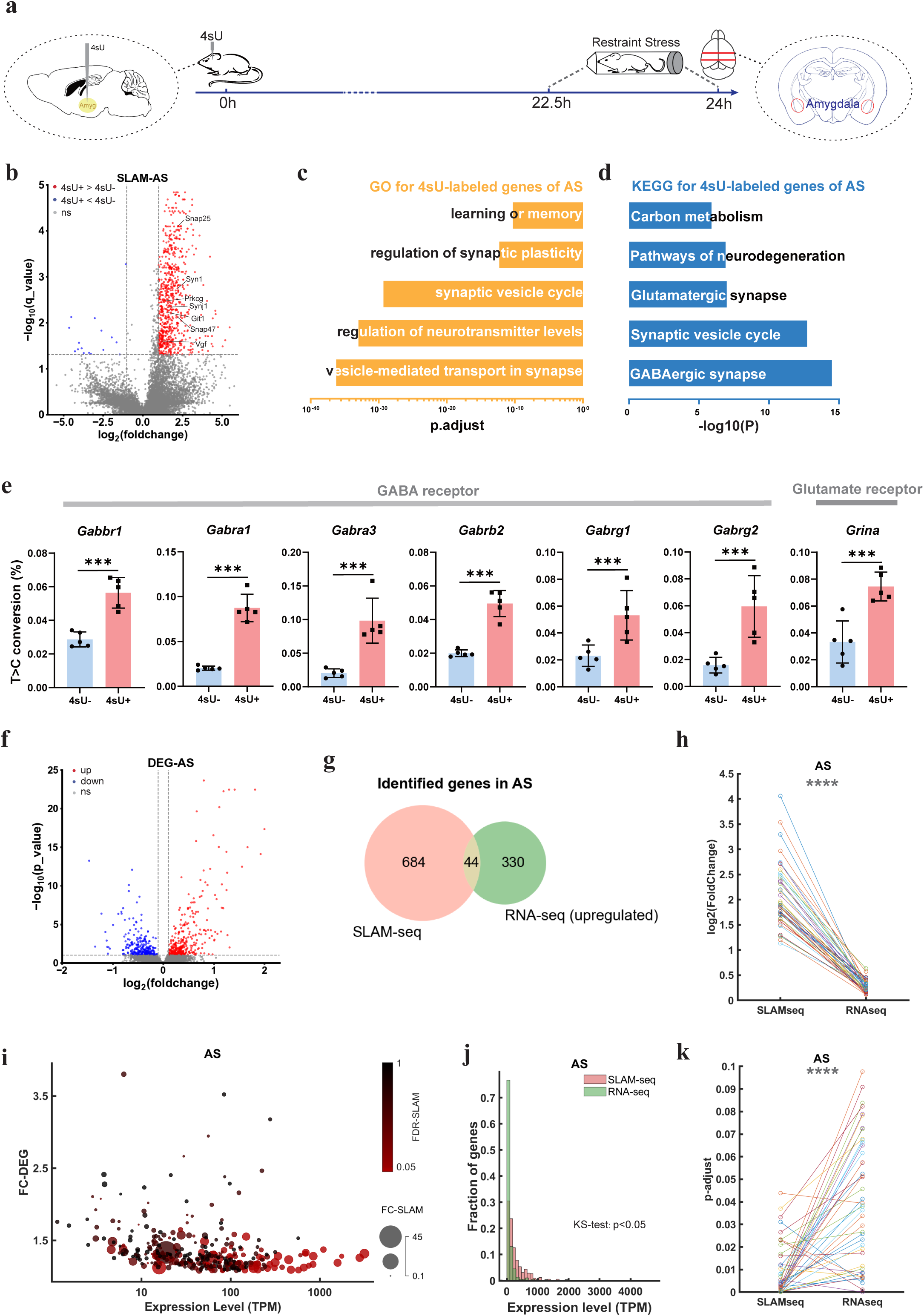
Transcriptome dynamics of mouse amygdala under acute stress. **(a)** Time line of acute stress experiment, including time points for 4sU injection and brain extraction. **(b)** Volcano plots showing T>C conversion rates of transcripts under AS, with or without 4sU injection. Red dots represent transcripts with significant higher T>C conversion rates in the 4sU+ group (FDR < 0.05 and log_2_FC > 1); blue dots represent transcripts with significant lower T>C conversion rates in the 4sU+ group (FDR < 0.05 and log_2_FC < -1). **(c)** GO term analysis for genes detected by SLAM-seq. **(d)** KEGG analysis for genes detected by SLAM-seq. **(e)** Fold changes of T>C conversion rates for GABA and glutamate receptors detected by SLAM-seq. Student’s t-test, *** P < 0.001. **(f)** Volcano plots showing differentially expressed genes (DEGs) under AS compared to BL. Red dots represent significantly AS-upregulated genes (FDR < 0.1 and log_2_FC > 0.1); blue dots represent significantly AS-downregulated genes (FDR<0.1 and log_2_FC < -0.1). ns, no significant difference; up, significantly upregulated; down, significantly downregulated. **(g)** Venn diagram showing intersection of AS-specific genes identified by SLAM-seq (red dots in **b**) and RNA-seq (red dots in **f**). **(h)** Fold changes of T>C conversion rates for 4sU+ vs. 4sU-(SLAM), and of expression levels for AS vs. BL (DEG) of all overlapped genes in (**g**). Paired t-test, *** P < 0.001. **(i)** Correlation between FC-DEG and expression levels of AS-upregulated genes in **3g**, with the color and size of each data point denoting the significance level (FDR-SLAM) and fold change (FC-SLAM) of each AS-upregulated gene. **(j)** Distribution for expression levels of genes detected by SLAM-seq and RNA-seq under AS. Two-sample Kolmogorov-Smirnov test, p < 0.05 **(k)** Significance levels of T>C conversion rates for 4sU+ vs. 4sU-(SLAM), and of expression levels for AS vs. BL (DEG) of all overlapped genes in (**g**). Paired t-test, *** P < 0.001.

By comparing the T>C conversion rates of genes between the 4sU and DMSO groups under AS, we identified 728 differentially 4sU-labeled genes with significantly higher T>C conversion rates in the 4sU group (FDR<0.05, log_2_FC>1, **Fig. 3b**. FDR: false discovery rate. FC: fold change). A gene ontology (GO) analysis of the differentially 4sU-labeled genes in the AS samples indicated enrichment for terms including “vesicle-mediated transport in synapse”, “regulation of neurotransmitter levels”, and “synaptic vesicle cycle”, all of which are closely associated with synaptic vesicle functions (**Fig. 3c, Supp.** Fig. 4b). Among these are genes tightly linked to synaptic vesicle release, including *Git1, Syn1, Synj1* [37–39] (**Fig. 3b**). GO analysis also revealed enrichment for terms such as “regulation of synaptic plasticity” and “learning or memory”, and related genes included *Vgf, Prkcg, Snap25, Snap47* [40–42] (**Fig. 3b, 3c**).

A KEGG pathway analysis showed that the most significant pathways include “glutamatergic synapse” and “GABAergic synapse”, suggesting that AS had a strong influence on both excitatory and inhibitory synaptic functions (**Fig. 3d**). Comparing genes identified under BL condition with genes identified under AS, we found only one GABA receptor gene (*Gabra1*) among the differentially 4sU-labeled genes under BL, in contrast to 6 GABA receptor genes under AS (*Gabbr1, Gabra1, Gabra3, Gabrb2, Gabrg1, Gabrg2*), representing a higher transcription rate for GABAergic receptors in the amygdala after acute stress. Conversely, only one glutamate receptor gene (*Grina*) was detected under AS, and none under BL (**Fig. 3e**). Consistent with previous studies linking amygdala GABAergic inhibitory control to stress response [14, 43], our results suggested a strong influence on GABAergic neurons in the amygdala by acute stress, in the form of quickly elevating GABA receptor expression.

### Sensitivity of SLAM-seq vs. RNA-seq

Neurons require high abundance of transcripts for maintaining normal functions, while also exhibit highly dynamic activity-induced transcription, making it hard to detect the subtle changes by evaluating the total RNA amount using classic RNA-seq [22]. Indeed, none of the differentially 4sU-labeled GABA receptor and glutamate receptor genes identified by SLAM-seq were detected using the differential gene expression analysis, even with a lenient standard (FDR<0.1, Log_2_FC>0.1, **Fig. 3f, Supp.** Fig. 4c). In total, 374 significantly upregulated and 279 downregulated genes were identified using DESeq2 by comparing RNA read counts between AS and BL conditions. 44 overlapping genes were identified by intersecting these AS-upregulated genes with differentially 4sU-labeled genes under AS (**Fig. 3g**). These genes include known stress-related genes such as *Cldn11, Vamp1, Glud1* and *Etnppl* [44–47]. For all of the overlapped genes, fold change is bigger for SLAM-seq analysis than RNA-seq (**Fig. 3h**).

Further investigation revealed that the overlapped genes detected by SLAM-seq and RNA-seq had relatively high expression levels (**Fig. 3i, Supp.** Fig. 4d). In fact, the genes with very high expression levels that were marginally upregulated as determined by DESeq2 (Log_2_FC <0.5) were detected by SLAM-seq (red data points in **Fig. 3i**) with bigger fold changes and higher significance levels (**Fig. 3h, k**). In addition, genes identified by SLAM-seq had higher expression levels than genes detected by RNA-seq (**Fig. 3j**, Kolmogorov-Smirnov test, p < 0.05). These results suggest that compared to RNA-seq, SLAM-seq is more sensitive to genes with high expression levels, and is therefore useful for detecting the increased active transcription of genes that can be masked by the high abundance of transcripts required for normal functioning. This is of particular importance for neurons, given the high basal expression level of many genes [22].

### Transcriptome dynamics of amygdala under chronic stress

Chronic stress (CS) poses prolonged impact in the brain [48]. For chronic stress experiments, we subjected mice to 7 days of restraint stress, 2 h each day [49, 50] (Methods, **Fig. 4a**). To examine the long-lasting transcriptomic changes under CS, we extracted the amygdala at 10 h after the final restraint stress session on day 7. We reasoned that by shifting 10 h, we reduced the possibility of detecting acute changes induced by the last session of restraint stress. 4sU was injected into the amygdala 24 h before brain extraction, same as BL and AS experiments. By comparing T>C conversion rates between 4sU and DMSO injected groups, we identified 815 differentially 4sU-labeled genes under CS (**Fig. 4b, Supp.** Fig. 5a). We performed a GO analysis for these genes, and found that in addition to terms associated with neuronal functions such as “synaptic vesicle cycle” and “neurotransmitter secretion”, there is also enrichment for terms closely linked to glial cells, especially oligodendrocytes and microglia (**Fig. 4c**). These terms include “myelination”, “ensheathment of neurons” and “axon ensheathment”, which are tightly linked to the function of oligodendrocytes, as well as “autophagy” and “autophagosome”, which are associated with microglia (**Supp.** Fig. 5b). Among these genes, *Mag* and *Dag1* are associated with myelination, and *Rragb* and *Usp36* are associated with autophagy [51].

**Fig. 4:**
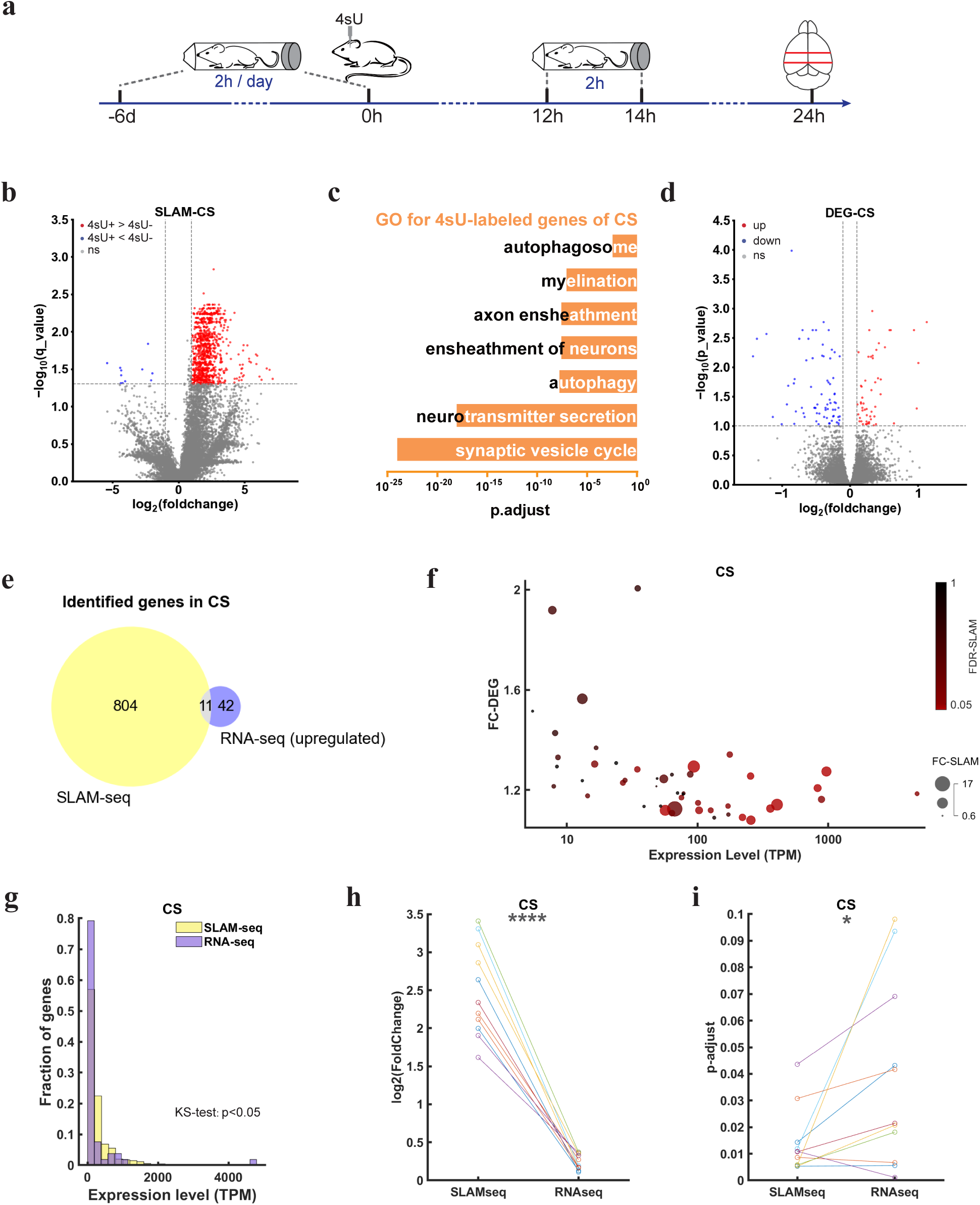
Transcriptome dynamics of mouse amygdala under chronic stress. **(a)** Time line of chronic stress experiment, including time points for 4sU injection and brain extraction. **(b)** Volcano plots showing T>C conversion rates of transcripts under CS, with or without 4sU injection. Red dots represent transcripts with significant higher T>C conversion rates in the 4sU+ group (FDR < 0.05 and log_2_FC > 1); blue dots represent transcripts with significant lower T>C conversion rates in the 4sU+ group (FDR < 0.05 and log_2_FC < -1). **(c)** GO term analysis for genes detected by SLAM-seq. **(d)** Volcano plots showing differentially expressed genes (DEGs) under CS compared to BL. Red dots represent significantly AS-upregulated genes (FDR < 0.1 and log_2_FC > 0.1); blue dots represent significantly AS-downregulated genes (FDR<0.1 and log_2_FC < -0.1). **(e)** Venn diagram showing intersection of AS-specific genes identified by SLAM-seq (red dots in **b**) and RNA-seq (red dots in **d**). **(f)** Correlation between FC-DEG and expression levels of AS-upregulated genes in **e**, with the color and size of each data point denoting the significance level (FDR-SLAM) and fold change (FC-SLAM) of each CS-upregulated gene. **(g)** Distribution for expression levels of genes detected by SLAM-seq and RNA-seq under CS. Two-sample Kolmogorov-Smirnov test, p < 0.05. **(h)** Fold changes of T>C conversion rates for 4sU+ vs. 4sU-(SLAM), and of expression levels for AS vs. BL (DEG) of all overlapped genes in (**e**). Paired t-test, *** P < 0.001. **(i)** Significance levels of T>C conversion rates for 4sU+ vs. 4sU-(SLAM), and of expression levels for AS vs. BL (DEG) of all overlapped genes in (**e**). Paired t-test, * P < 0.05.

Regarding sensitivity for SLAM-seq vs. RNA-seq, we noted a similar trend for CS data to AS. DESeq2 analysis detected 53 significantly up-regulated and 11 down-regulated genes for CS (FDR<0.1, Log_2_FC>0.1, **Fig. 4d**). Intersection of the differentially 4sU-labeled genes identified by SLAM-seq and upregulated genes identified by RNA-seq revealed 11 overlapping genes (**Fig. 4e**), all with high expression levels (**Fig. 4f, Supp.** Fig. 5c). Expression of six out of the 11 overlapped genes is affected by chronic stress (*Esrrg, Gabarapl2, Cct8, Vamp1, Nrsn1, Cplx1*) [47, 52–54]. These genes all show bigger fold changes and 9 out of 11 genes also show higher significance levels for SLAM-seq than for RNA-seq (**Fig. 4h, 4i**). Similar to AS, genes identified by SLAM-seq in CS also had higher expression levels (**Fig. 4g**, Kolmogorov-Smirnov test, p < 0.05), again demonstrating the sensitivity of SLAM-seq for detecting actively transcribed genes with high abundance. The GO analysis of the genes identified by SLAM-seq indicated enrichment for terms closely linked to glial functions, but these terms did not show in GO analysis of genes identified by RNA-seq (**Supp.** Fig. 6). Thus, the low-magnitude yet persistent transcriptome changes associated with CS are more likely to be captured by SLAM-seq.

### Comparison of differentially 4sU-labeled genes among BL, AS and CS conditions

By comparing the differentially 4sU-labeled genes under BL, AS and CS conditions, we found that a majority of genes (363 out of 413) labeled under BL were also identified in either AS or CS conditions, including known housekeeping genes such as *Sdha, Ywhaz*, *Gfap* [55] (**Fig. 5a**). These genes were actively transcribed regardless of stress conditions, indicating their important roles in maintaining normal cellular functions. In contrast to BL, we identified many more condition-specific genes after stress treatments, namely, 224 AS-specific genes and 333 CS-specific genes (**Fig. 5a**). Among the 172 AS-CS overlapped genes, 29 were associated with synaptic transmission, synaptic vesicle cycle and synaptic regulations (**Fig. 5a**). Thus, both acute and chronic stress strongly affected synaptic functions in amygdala neurons. Notably, compared to CS, AS triggered active transcription of a higher proportion of synapse-related genes, among which were genes associated with protein localization to postsynaptic membrane (*Git1, Mapk10, Stx1b, Kif1a*) [39, 56, 57], highlighting immediate postsynaptic plasticity caused by acute stress (**Fig. 5b**).

**Fig. 5:**
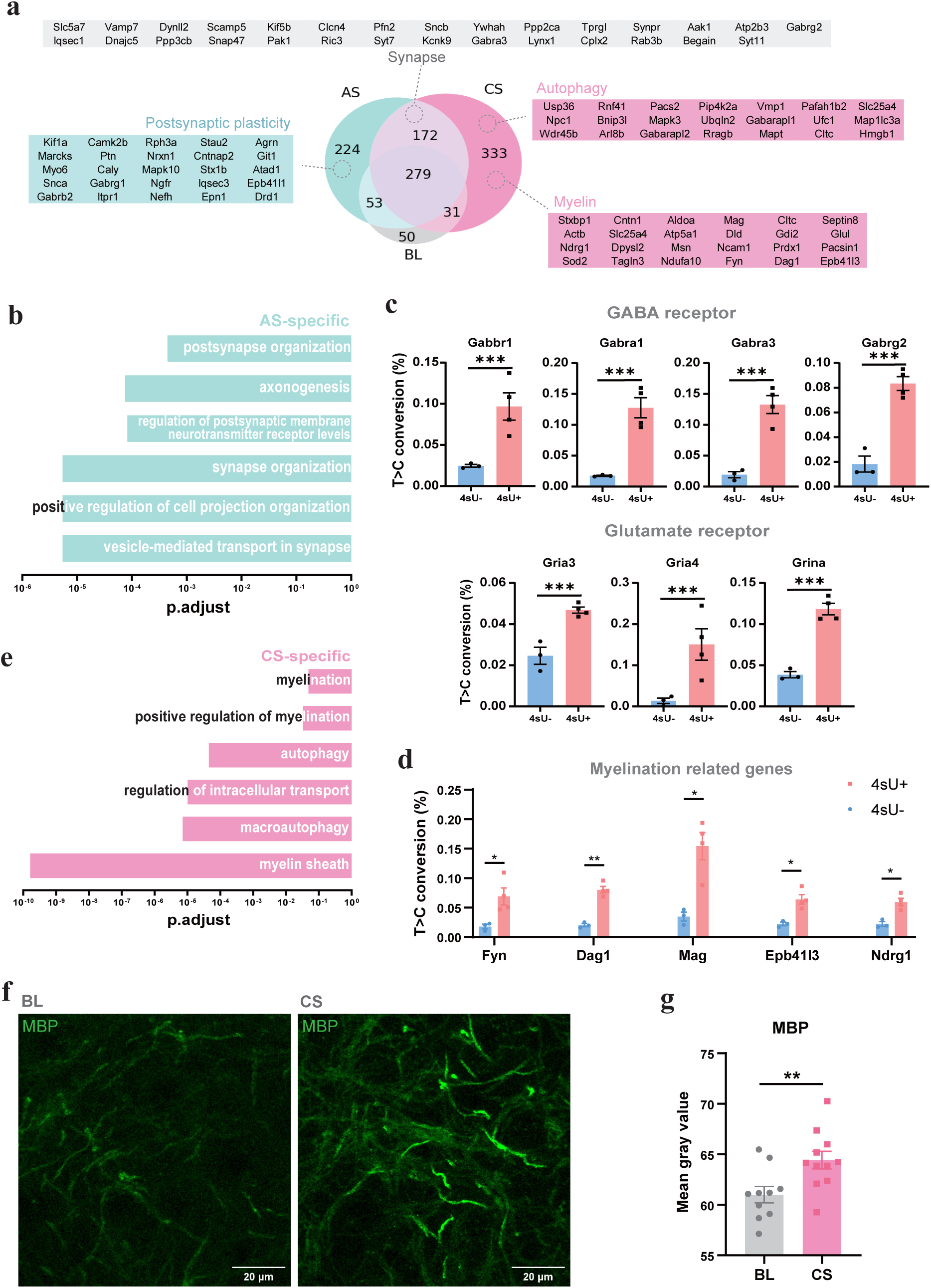
Comparison of AS and CS-associated genes and validation of SLAM-seq results. (a) Venn diagram showing intersections of genes identified by SLAM-seq under BL, AS and CS conditions. Genes associated with synapse, postsynaptic plasticity, autophagy and myelination are listed. (b) GO term analysis for AS-specific genes detected by SLAM-seq. (c) Fold changes of T>C conversion rates for GABA and glutamate receptors detected by SLAM-seq under CS. Student’s t-test, *** P < 0.001. (d) Fold changes of T>C conversion rates for myelination related genes detected by SLAM-seq under CS. Student’s t-test, *** P < 0.001. (e) GO term analysis for CS-specific genes detected by SLAM-seq. (f) Example images showing immunostaining of myelin basic protein (MBP) in the amygdala of control (BL) and chronically stressed (CS) mice. Scale bar, 20 μm. (g) Increased MBP signal intensity in CS mice (n=5) compared to BL mice (n=5), measured by mean gray value. Student’s t-test, ** p<0.01.

As described in an earlier section, we detected 6 GABA receptor genes (*Gabbr1, Gabra1, Gabra3, Gabrb2, Gabrg1, Gabrg2*) and 1 glutamate receptor gene (*Grina*) under AS, representing an imminent effect of AS on GABAergic inhibitory control of the amygdala. In comparison, we found 4 GABA receptor genes (*Gabbr1, Gabra1, Gabra3, Gabrg2*) and 3 glutamate receptor genes (*Gria3, Gria4, Grina*) among the differentially 4sU-labeled genes under CS (**Fig. 5c**). The increase in excitatory receptor transcription and relatively lower transcription of GABAergic receptors might underlie hyperactivity of the amygdala [13, 14], caused by stress-induced removal of GABAergic inhibitory control.

### Validation of SLAM-seq results by quantitative immunofluoresence

Recent studies suggested that elevated neuronal and axonal activity could trigger myelination [20, 21], a phenomenon known as “activity-dependent myelination”. In line with these results, CS-specific genes detected by SLAM-seq included a large proportion of genes related to myelination, including *Fyn, Dag1, Mag, Epb41l3,* and *Ndrg1* (**Fig. 5d**), and the enriched GO terms included “myelination”, “positive regulation of myelination”, and “myelin sheath” (**Fig. 5e**). These genes were not detected by RNA-seq (**Supp.** Fig. 7), possibly due to their high baseline expression levels. To examine whether the increased transcription of myelination related genes after CS is indicative of increased myelination in the amygdala, we stained brain slices of chronically stressed mice and control mice with a myelin marker, the myelin basic protein (MBP), and compared the immunofluorescence signal intensity between the two groups (**Fig. 5f**). We found that MBP signal is indeed significantly stronger in the chronically stressed amygdala than the control (p<0.01, Student’s t-test, **Fig. 5g**), validating the SLAM-seq results that chronic stress led to increased myelination in the mouse amygdala.

## Discussion

By directly injecting 4-thiouridine (4sU) into the amygdala followed by SLAM-seq, we were able to identify newly transcribed genes under acute and chronic stress. By cross-referencing these genes with known cell type-specific marker genes, we found that the injected 4sU was integrated into the transcription of major subtypes of neurons and glia cells in the amygdala. Analyses revealed that acute stress resulted in increased transcription of GABA receptors in neurons, while chronic stress had a stronger impact on transcription of glutamate receptors and genes associated with myelination, both in line with heightened activity of the amygdala [14]. Furthermore, we observed that SLAM-seq exhibited greater sensitivity to transcriptional changes in genes with high expression levels compared to RNA-seq. Thus, by applying SLAM-seq to the amygdala, we generated a rich dataset which can be used in future studies to dissect the functions of genes related to stress responses.

Neurons have highly specialized morphology, including axons and dendrites that can span long distances, with functional subcellular compartments such as dendritic spines and axonal boutons [58]. The transcriptional networks in neurons are dynamically modulated by synaptic activities, through coupling of a small subset of active synapses to the nucleus [59, 60]. Given the abundance of gene transcripts for maintaining fundamental neuronal functions such as action potential firing [19], the transcriptional modulation that lead to activation of relevant gene expression may not alter the total mRNA of these genes [22]. Therefore, the most common method for gene expression profiling, RNA-seq, may not be suited for accessing the highly dynamic transcriptional changes under fast-varying physiological, behavioral or pathological conditions.

SLAM-seq, a metabolic RNA labeling method for identifying newly synthesized RNA molecules, is an alternative approach to gain insights into the transcriptional regulation regardless of the steady-state abundance of genes [24–26]. A previously published method, SLAM-ITseq, extended SLAM-seq to mouse tissues by crossing Cre-dependent UPRT mice with Cre lines [27]. This method requires usage of transgenic animals, and the cell types that can be investigated are restricted by available Cre lines. Instead of intravenously injecting 4-thiouracil and relying on Cre-dependent UPRT expression for cell-specific labeling, we used stereotactic 4sU injection to target all cell types in selected brain regions. Our data showed that 4sU can be effectively imported into major cell types [28], and incorporated into the transcription, without causing detectable tissue damage or cell death. This is a simplified method that can be applied to any brain region in mice and possibly other species using stereotaxic injection, without the need for transgenic animals.

We used this method to investigate the RNA dynamics of a key brain region involved in stress conditions, the amygdala, under acute and chronic restraint stress. We found partial overlap in genes detected by SLAM-seq and RNA-seq, and over half of the overlapped up-regulated genes under CS condition had been implicated in chronic stress by previous studies [52–54, 61–64]. Notably, for all these genes, SLAM-seq showed higher sensitivity than RNA-seq, shown by higher fold changes and significance levels. The upregulated genes with high abundance were detected by SLAM-seq with high significance level, while barely meets the lenient criteria in RNA-seq. It supports the notion that for genes with high expression rate, physiological condition (restraint stress) may not cause a noticeable change in the total amount of mRNA for relevant genes. One such example is the ensemble of genes related to myelination. The high abundance of these genes precluded detection by RNA-seq following CS treatment, although immunostaining of myelin-specific protein revealed increased myelination, consistent with the SLAM-seq results. Therefore, SLAM-seq serves as a better method for screening the dynamics of highly expressed genes.

Competition and cooperation between excitation and inhibition is crucial for balancing neural activity[65, 66]. Using SLAM-seq, we identified 6 GABA receptors under AS, compared to 1 under BL and 4 under CS, suggesting an imminent inhibition drive induced by AS. Conversely, 3 glutamate receptors were identified under CS, while only 1 under AS and 0 under BL, suggesting an increase in elevated activity of the amygdala. Furthermore, CS also induced transcription of many genes associated with myelination. Myelination has been shown to be initiated by neuronal activities [20, 21], and can in turn facilitate signal transduction in neurons [51, 67]. These results are consistent with previous findings that stress could cause hyperactivity of the amygdala [14, 68, 69]. Therefore, our findings indicated that acute and chronic stress could differentially modulate the transcription level of GABA and glutamate receptors as well as proteins constituting the myelin sheath, and in turn influence the excitation-inhibition balance of the amygdala neurons.

We provided a rich dataset of molecular responses of amygdala to acute and chronic stress, and confirmed that CS can induce increased myelination through immunostaining. Future studies will be essential for understanding the functions of other genes identified in this study.

## Acknowledgements

We thank Drs. Margaret Ho and Dan-Qian Liu for comments on the manuscript. We thank Dr. Hao Wu and Ms. Liang He for assistance in data analysis and experiments. We thank the animal facility of ShanghaiTech for their excellent care of mice. This work was supported by grants from the Ministry of Science and Technology of China (2022ZD0204900), Central Guidance on Local Science and Technology Development Fund (YDZX20233100001002), and ShanghaiTech University Startup Fund to Y.Y.

## Materials and methods

### Animals

C57BL/6 mice were purchased from Slac Laboratory Animals. Mice were housed and bred in a 12 h light-dark cycle (7 am - 7 pm light) in the animal facility of ShanghaiTech University. Eight- to ten-week-old male mice were used for the experiments. All procedures were approved by the Animal Committee of ShanghaiTech University.

### 4sU Injection

The 4sU powder was dissolved in dimethyl sulfoxide (DMSO) to reach a concentration of 1 M, and the solution was aliquoted, frozen, and stored at -20°C. Before injection, the solution was 1:10 diluted with phosphate-buffered saline (PBS). For injection, mice were anesthetized with isoflurane (1.5-2 %) and positioned in a stereotaxic frame (RWD Life Science Co.). Body temperature was maintained at 37 °C using a heating pad. 4sU solution (1 μL) were injected using a glass micropipette with a tip diameter of 15–20 μm, through a small skull opening (<0.5 mm2), with a microinjector (Nanoject III), at a rate of 1 nL/s. In baseline, acute stress and chronic stress experiments, 4sU was injected into bilateral amygdala 24 h before brain extraction.

### TUNEL Staining

The TUNEL assay was performed following the one-step method outlined in the Promega TUNEL Cell Apoptosis Detection Kit. Mice were intracranially injected with 1 μL of 100 mM 4sU into the auditory cortex (M/L=-4.4 mm, A/P=-2.2 mm, D/V=-1.1 mm) or amygdala (M/L=±3.25 mm, A/P=-1.1 mm, D/V=-3.95 mm), as described above. After 24 h, mice were deeply anesthetized and transcardially perfused with PBS and paraformaldehyde (PFA), followed by brain extraction. The extracted brains were postfixed in PFA at 4°C for 12 hours and embedded in agarose for vibratome sectioning at 70 μm per slice. Brain slices were collected for TUNEL staining, in which terminal deoxynucleotidyl transferase (TdT) labels DNA strand breaks with fluorophore-labeled nucleotides. Stained brain slices were mounted on glass slides and imaged using confocal fluorescence microscopy (Leica).

### Tissue RNA Extraction

Mice were deeply anesthetized and transcardially perfused with cold PBS. Each mouse was then quickly decapitated, and the head was immersed in ice-cold PBS for rapid cooling for 10 seconds. The mouse brain was then carefully extracted using forceps, placed in a pre-chilled mouse brain mold (RWD Life Science Inc.), and a 2 mm-thick brain tissue block containing the target region (auditory cortex or amygdala) was sliced using a razor blade. The brain slice was then placed on a 6 mm petri dish and dropped into liquid nitrogen briefly for 7-8 s. Then, the auditory cortex or amygdala was cut out using a 15 G needle, and promptly transferred to cryotubes for rapid freezing in liquid nitrogen, and subsequent storage at -80°C.

RNA extraction from mouse brain tissue was performed using the Trizol method. The frozen tissue in cryotubes was thawed, and 1 mL of Trizol (MagZol Reagent R4801-02) was added to each tube. Tissue homogenization was performed using an electric homogenizer, with a 10-second pause every 10 seconds to prevent overheating. The mixture was then left at room temperature for 5 minutes for tissue digestion, and the liquid was transferred to 1.5 mL centrifuge tubes. To each tube, 200 μL of chloroform (one-fifth of the Trizol volume) was added, and after thorough vortex mixing, the mixture was left at room temperature for 15 minutes. The mixture was then centrifuged at 12,000 rpm for 15 minutes at 4°C. Then, the supernatant containing RNA was carefully aspirated into new tubes, an equal volume of pre-cooled isopropanol was added, along with 1/100 volume of 10 mM DTT (final concentration 0.1 mM) and 1 μL glycogen (20 mg/mL). After thorough mixing, the mixture was left at -20°C for 1 hour for RNA precipitation. Subsequently, it was centrifuged at 12,000 rpm for 30 minutes at 4°C, resulting in a white gel-like RNA substance at the bottom. The RNA pellet was washed once with pre-cooled 75% ethanol (containing 5 μL 10 mM DTT), and after removing the supernatant, the RNA pellet was air-dried for approximately 3-5 minutes. Finally, the RNA was dissolved in 20 μL RNase-free water, and the concentration was determined using Nanodrop 2000.

### Dot Blot Assay

Four RNA samples with different 4sU injection conditions were used in this experiment: 1) RNA extracted from cortex with no 4sU injection, 2) RNA extracted from cortex dissected 2 h after 4sU injection, 3) RNA extracted from cortex dissected 8 h after 4sU injection, 4) RNA extracted from cortex dissected 24 h after 4sU injection. Positive control was 5′-biotin labeled DNA oligo. Positive control and the extracted RNA samples (1 μg for each sample) were used for dot blot essay following the standard protocol for Chemiluminescent Biotin-labeled Nucleic Acid Detection Kit (Beyotime, D3308), using EZ-Link™ HPDP-Biotin (Thermo Scientific, 21341).

### Restraint Stress Experiment

Mice were acclimated in the animal facility for at least one week before the initiation of restraint stress experiments.

#### Acute stress group

Each mouse was carefully removed from its home cage and put in a Falcon 50 mL centrifuge tube, with 8 small ventilation holes drilled on the tube walls. Once the mouse entered the tube, the rear was sealed with a clean tissue, and the tube was capped with the lid, preventing forward or backward movements. Following the 1.5-hour restraint, the mouse was perfused and the brain was extracted.

#### Chronic stress group

Each mouse was carefully removed from its home cage and put in a 50 mL tube, as described in the acute stress experiment. Daily restraint sessions lasting 2 hours (from 10:00 am to 12:00 pm) were conducted for one week. On the 7^th^ day, 10 hours after the final restraint session ended, the mouse was perfused and the brain was extracted.

#### Baseline group

Mice did not undergo any restraint stress, and remained in their home cages until they were perfused.

### IAA treatment

Extracted RNA was reacted with IAA to alkylate the thiol group following [1]. For a 50 μL reaction: 2-3 μg RNA, 25 μL DMSO, 5 μL 100 mM iodoacetamide (IAA), and 5 μL 500 mM sodium phosphate (pH=8), and RNase-free water. IAA was freshly prepared and diluted with anhydrous ethanol. The prepared mixture was incubated at 50°C for 15 minutes, followed by the addition of 1 M dithiothreitol (DTT) to terminate the reaction. Then, 125 µL anhydrous ethanol, 1 µL glycogen (20 mg/mL), and 5 µL sodium acetate (3 M, pH=5.2) were added to the reaction system. After thorough mixing, the solution was left at -80°C for 30 minutes, followed by centrifugation at 12,000 rpm for 30 minutes at 4°C, resulting in a white gel-like RNA substance at the bottom. The RNA pellet was washed once with pre-cooled 75% ethanol. After removing the supernatant, the RNA pellet was air-dried for approximately 3-5 minutes, and dissolved in 20 μL RNase-free water.

### RNA sequencing

The IAA-methylated RNA was stored in a -80°C freezer, and library construction and sequencing were conducted by Suzhou Geneseeq Technology Inc. The libraries were sequenced using the Illumina NovaSeq for 150-base pair paired-end sequencing, generating approximately 6 GB of sequencing data for each RNA sample.

### Bioinformatics Analysis

#### Preprocessing of RNA-Seq Data

Raw data obtained from RNA-Seq was preprocessed using Cutadapt v3.1. Adapter sequences in paired-end sequencing were eliminated, and sequences shorter than 15 nucleotides were discarded. The preprocessed data was then aligned to the NCBI reference mouse genome (mm10) using Hisat2 v2.1.0. The resulting alignment data in SAM format was converted to BAM format using Samtools v1.9.

#### SLAM-Seq Analysis

The software SLAM-DUNK v0.2.4 (tneumann.github.io/slamdunk/), designed for SLAM-Seq analysis, was used to compute the T>C mutation counts for each sample [2]. Annotation data for 3’ UTR regions were obtained from the RefSeq and Ensembl databases. To identify significantly 4sU-labeled transcripts, a two-step filtering process was implemented. Firstly, genes with zero T read counts were excluded. Secondly, only genes with read counts > 5 in all samples were kept for further analyses. Statistical analysis was performed using the ibb R package (version 13.06), as described in [3]. To control for the false discovery rate (FDR), adjusted P-values were calculated using the Benjamini-Hochberg method. Transcripts with an FDR below 0.05 were considered significantly labeled.

#### Differential Gene Expression (DEG) Analysis

Quantification of gene expression was performed using FeatureCounts (v2.0.1). The Transcripts Per Million (TPM) values for each gene were calculated using the formula:

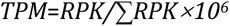

Here, RPK (Reads Per Kilobase) represents the reads per thousand base pairs, and ∑RPK is the sum of RPK values for all genes in the sample. Differential expression analysis was performed using the DESeq2 package in R 3.6.1. The standard DESeq2 workflow was followed for data normalization, variance estimation, and statistical modeling. Identification of significantly differentially expressed genes was based on the criterion of adjusted P-values < 0.05 or < 0.1.

### GO term enrichment analysis

List of genes was used as input for PANTHER 13.1 (pantherdb.org/) to perform ‘Statistical enrichment test’ with GO terms for biological processes, cellular components and molecular functions. The enriched GO terms obtained were further selected based on the P-values for visualization.

### Immunofluorescence

Chronic stressed mice were perfused 2 weeks after the final restraint session ended, with 0.01M PBS and 4% PFA. Control mice were perfused without any stress treatment. Brains were sliced (at 50 μm) using a vibratome (Leica VT1200S) after post-fixed for 5 d in 4% PFA at 4 °C. Brain slices including the amygdala were permeabilized with 0.4% Triton X-100 in PBS for 30 minutes, blocked with 5% bovine serum albumin (BSA) in PBS for 1 h at RT, and incubated with rat anti-MBP (1:200; MCA409S, Serotec) in 0.1% Triton X-100 and 1% BSA in PBS for 40 h at 4℃, then washed with PBS 3 × 10 minutes. Sections were incubated with DAPI (1:2500) and fluorophore-conjugated secondary antibodies (1:500; AB150165, abcam) in 0.1% Triton X-100 and 1% BSA in PBS for 2h at RT, then washed with PBS 3 × 10 minutes. Images were acquired at a Leica STELLARIS 8 FALCON, with a HC FLUOTAR L VISIR 25x/0.95 water objective; pinhole was set to 1 AU, pixel size to 113.55nm × 113.55nm and z-stack interval to 1 μm.

Image processing and quantification were performed using FIJI. For the quantitative immunofluorescence analysis, the mean grey value of each field of view was first thresholded for identification of myelin structures, and then calculated for 11 optical planes of acquired z-stacks (step 1 μm). For each mouse, the mean grey value is averaged across two sections containing the amygdala, one from each hemisphere.

