## Supplemental figures for "Transcriptome Dynamics in Mouse Amygdala under Acute and Chronic Stress Revealed by Thiol-labeled RNA Sequencing"

Supp. Fig. 1

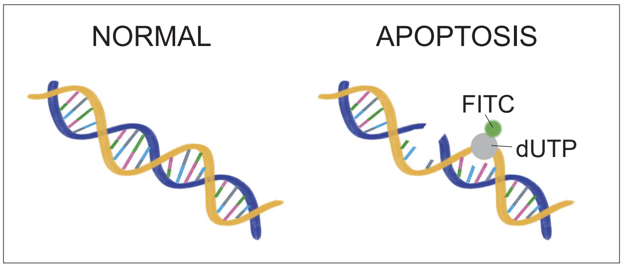


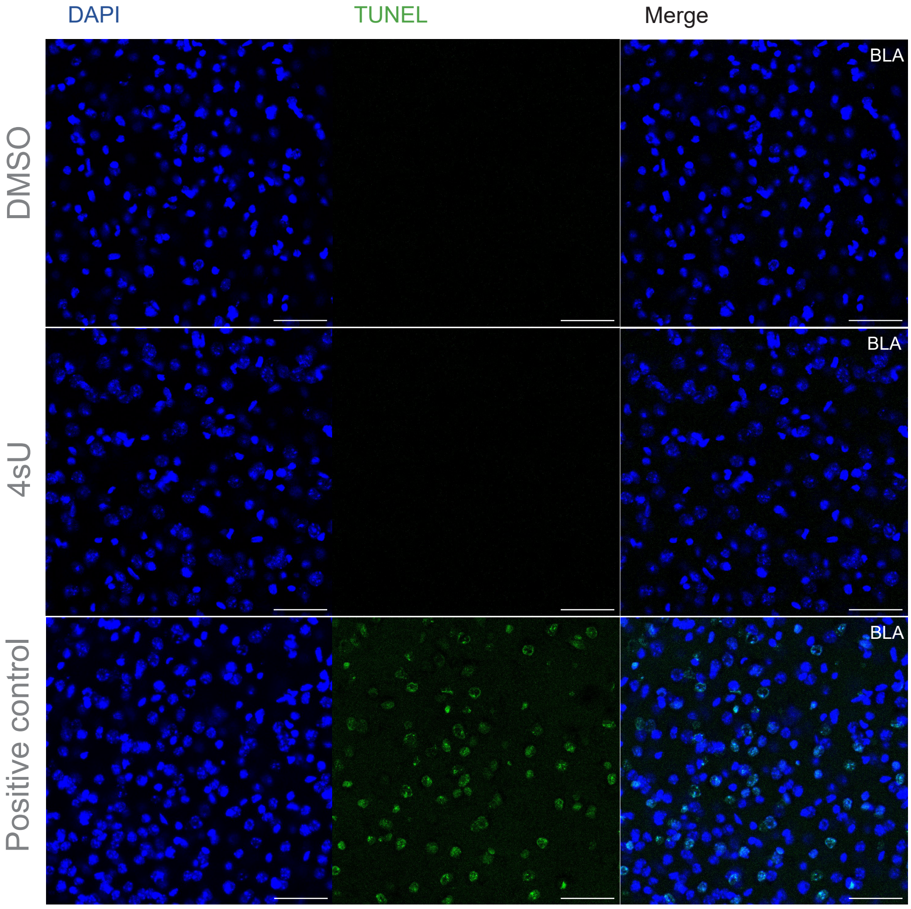


TUNEL staining of brain slices for positive control (top panels), and brain slices with DMSO (middle) and 4sU (bottom) injections in the amygdala. Green signals denoted dead cells. Scale bar = 50 µm.

Supp. Fig. 2

**a**

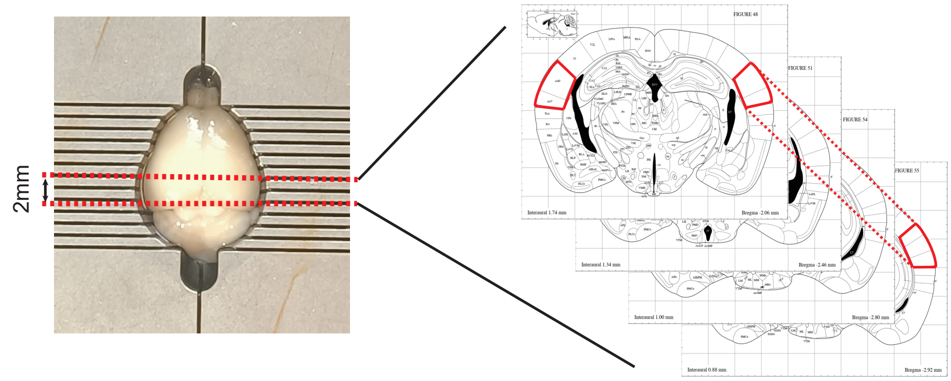


**b**

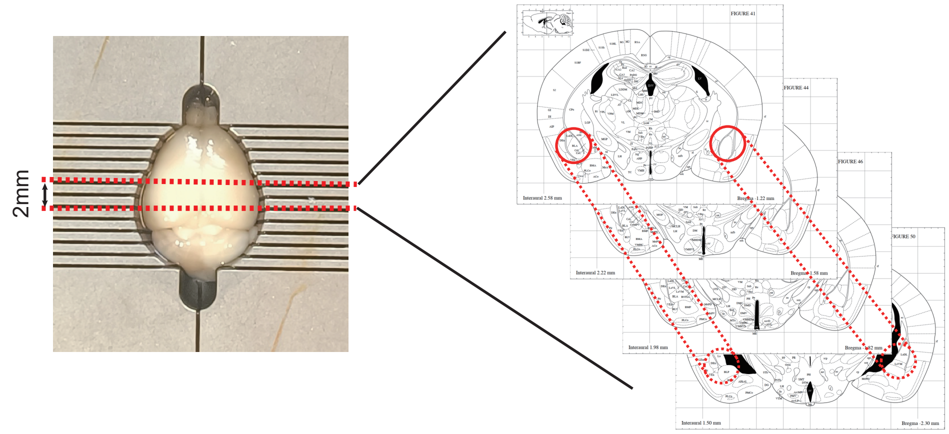


**c**

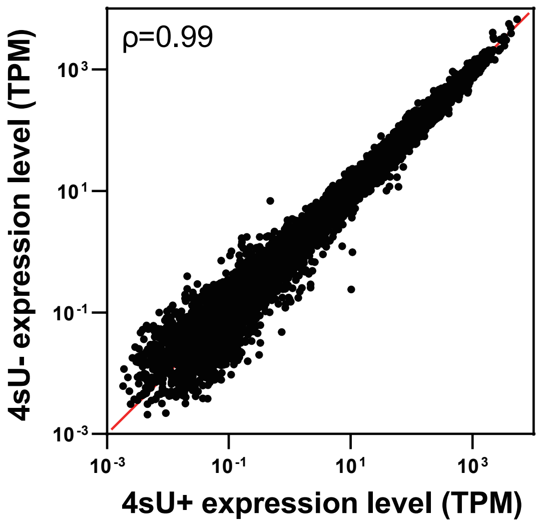


(a) Schematic showing position of auditory cortex tissue for RNA extraction.
(b) Schematic showing position of amygdala tissue for RNA extraction.
(c) Correlation of RNA levels extracted from DMSO and 4sU injected amygdala samples. ρ, spearman correlation coefficient.

Supp. Fig. 3

**a**

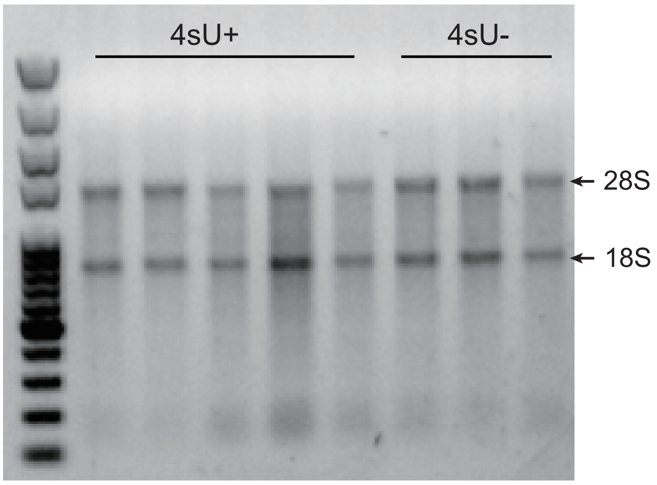


**b**

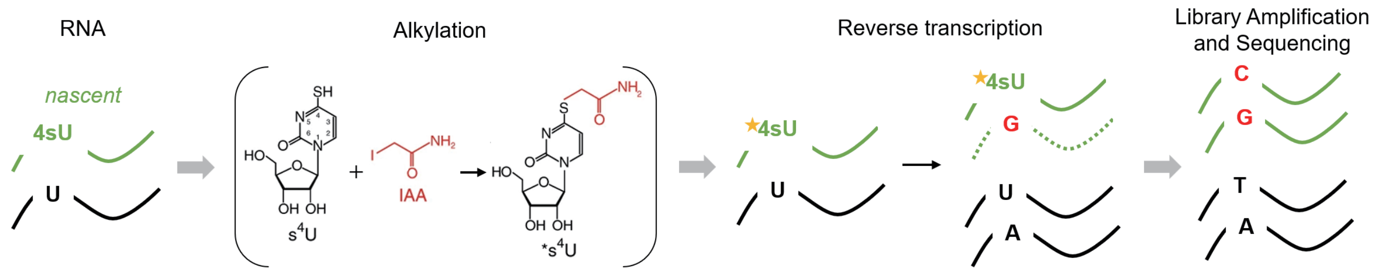


(a) Example RNA gel for 5 amygdala RNA samples injected with 4sU and 3 samples injected with DMSO after IAA treatment.
(b) Schematic showing IAA treatment and library preparation for SLAM-seq.

Supp. Fig. 4

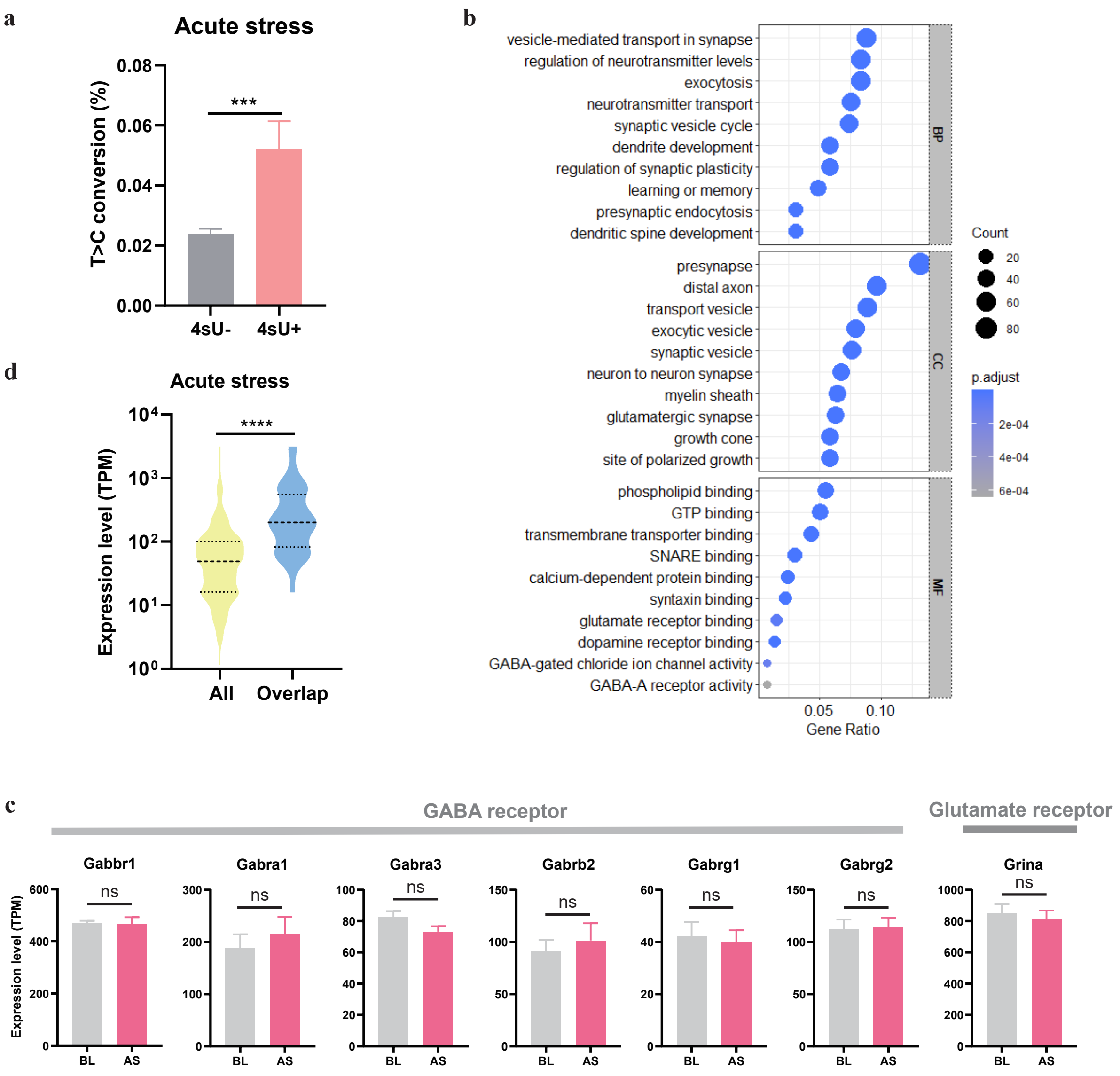


(a) T>C Conversion rates of total mRNA extracted from DMSO (n=5) and 4sU (n=5) injected amygdala under AS condition (two-tailed unpaired t-test, $p < 0.001$).
(b) GO term analysis for genes detected by SLAM-seq under AS, including BP, CC and MF.
(c) Gene expression levels for GABA and glutamate receptors, as shown in Fig. 3e.
(d) Violin plots showing gene expression levels for all upregulated-genes in Fig. 3i (All) and overlapped genes identified by SLAM-seq and RNA-seq (Overlap) under AS (two-tailed unpaired t-test, $p < 0.0001$).

Supp. Fig. 5

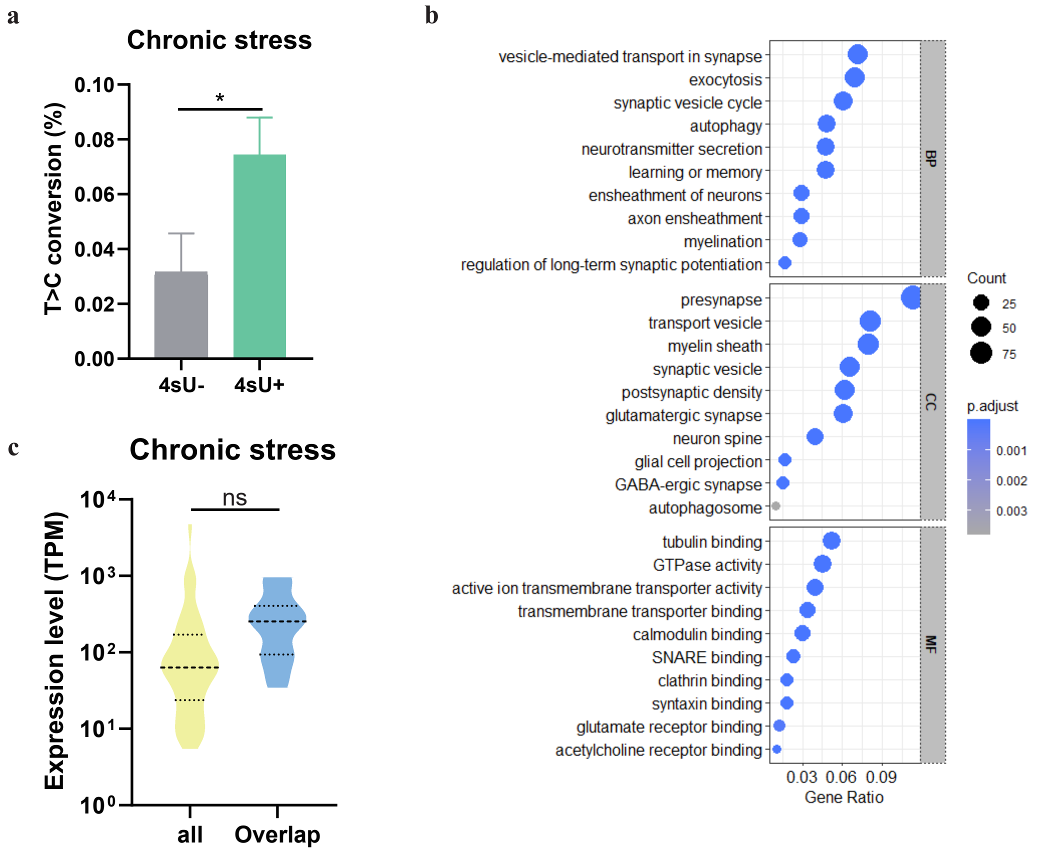


(a) T>C Conversion rates of total mRNA extracted from DMSO (n=4) and 4sU (n=3) injected amygdala under CS condition (two-tailed unpaired t-test, $p < 0.05$).
(b) GO term analysis for genes detected by SLAM-seq under CS, including BP, CC and MF.
(c) Violin plots showing gene expression levels for all upregulated-genes in Fig. 4f (All) and overlapped genes identified by SLAM-seq and RNA-seq (Overlap) under CS.

Supp. Fig. 6

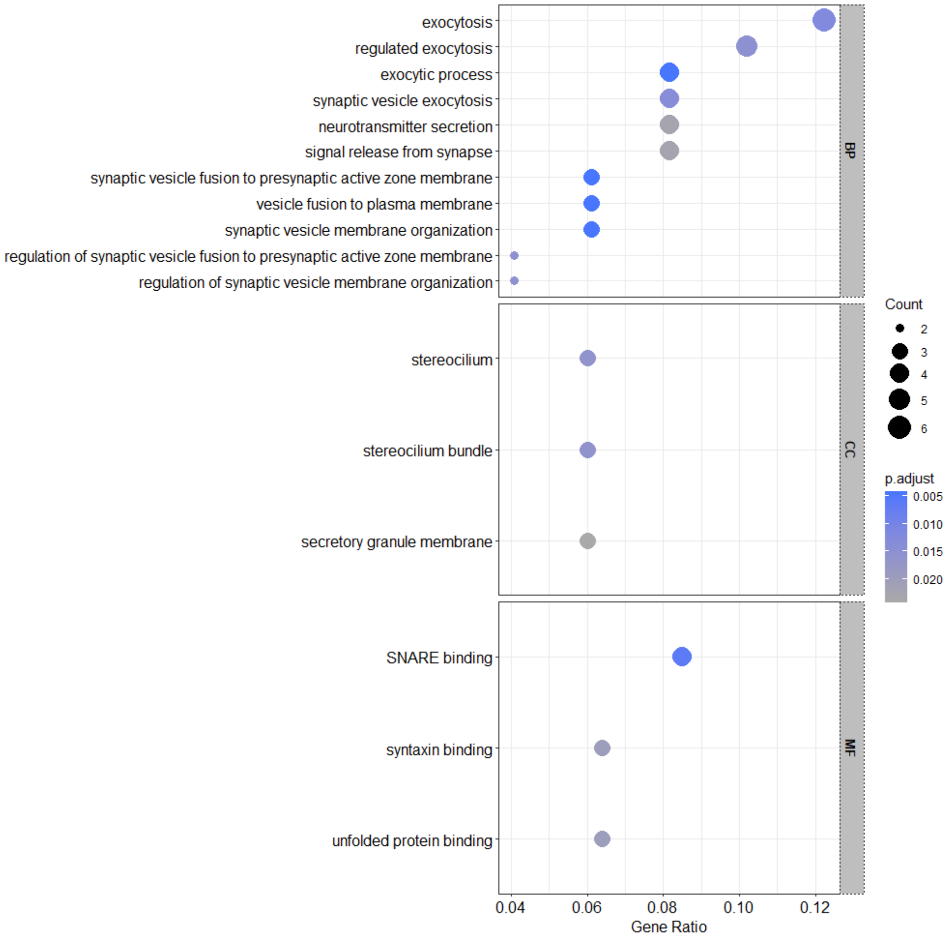


GO term analysis for AS-CS overlapped genes, including BP, CC and MF.

Supp. Fig. 7

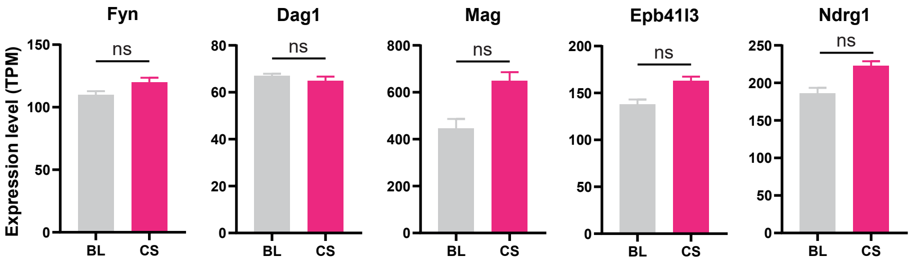


Expression levels for genes related to myelination, as shown in Fig. 5d.
